## Supplementary tables and figures for "A large scale, multiple genome comparison of acidophilic Archaea (pH ≤ 5.0) extends our understanding of oxidative stress responses in polyextreme environments": supplementary_figures.pdf

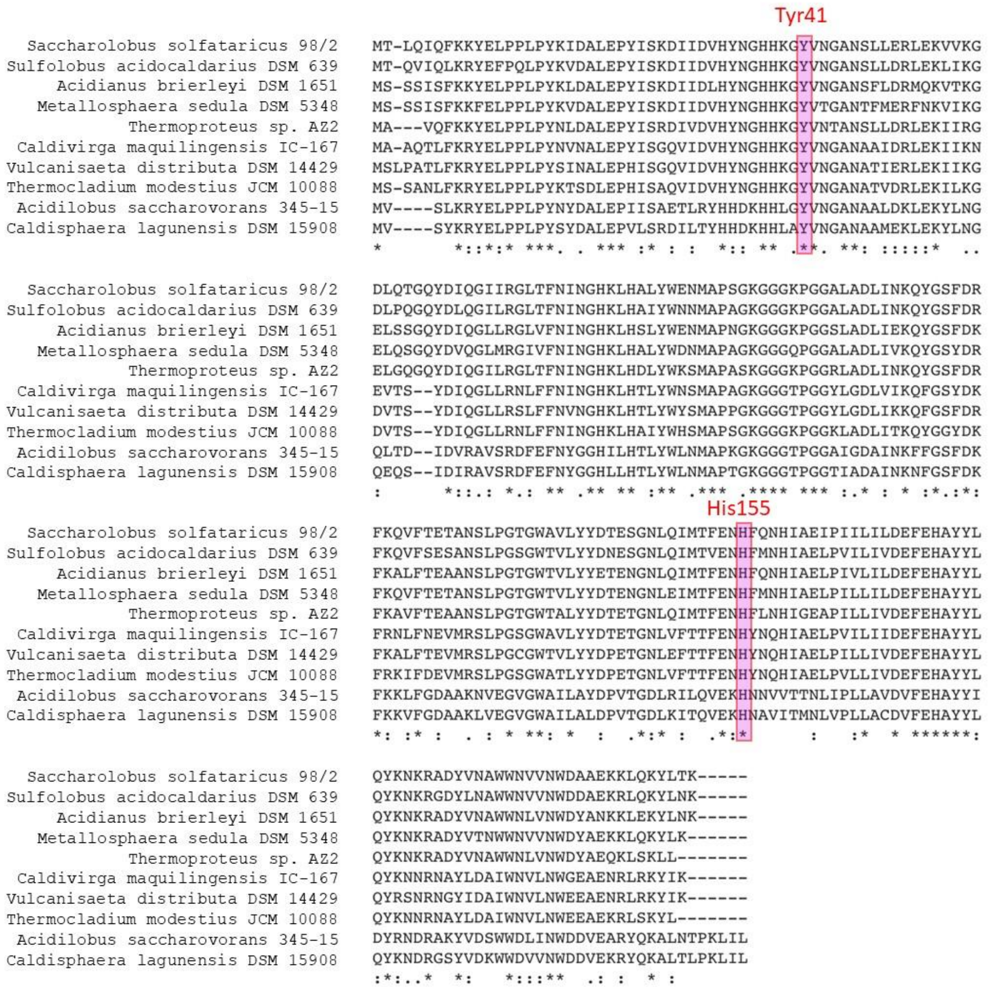


**Figure S1.** Multiple sequence alignment of Crenarchaeota Fe-SOD. In red are emphasized the conserved key amino acids Tyr41 and His155 through the different sequences.

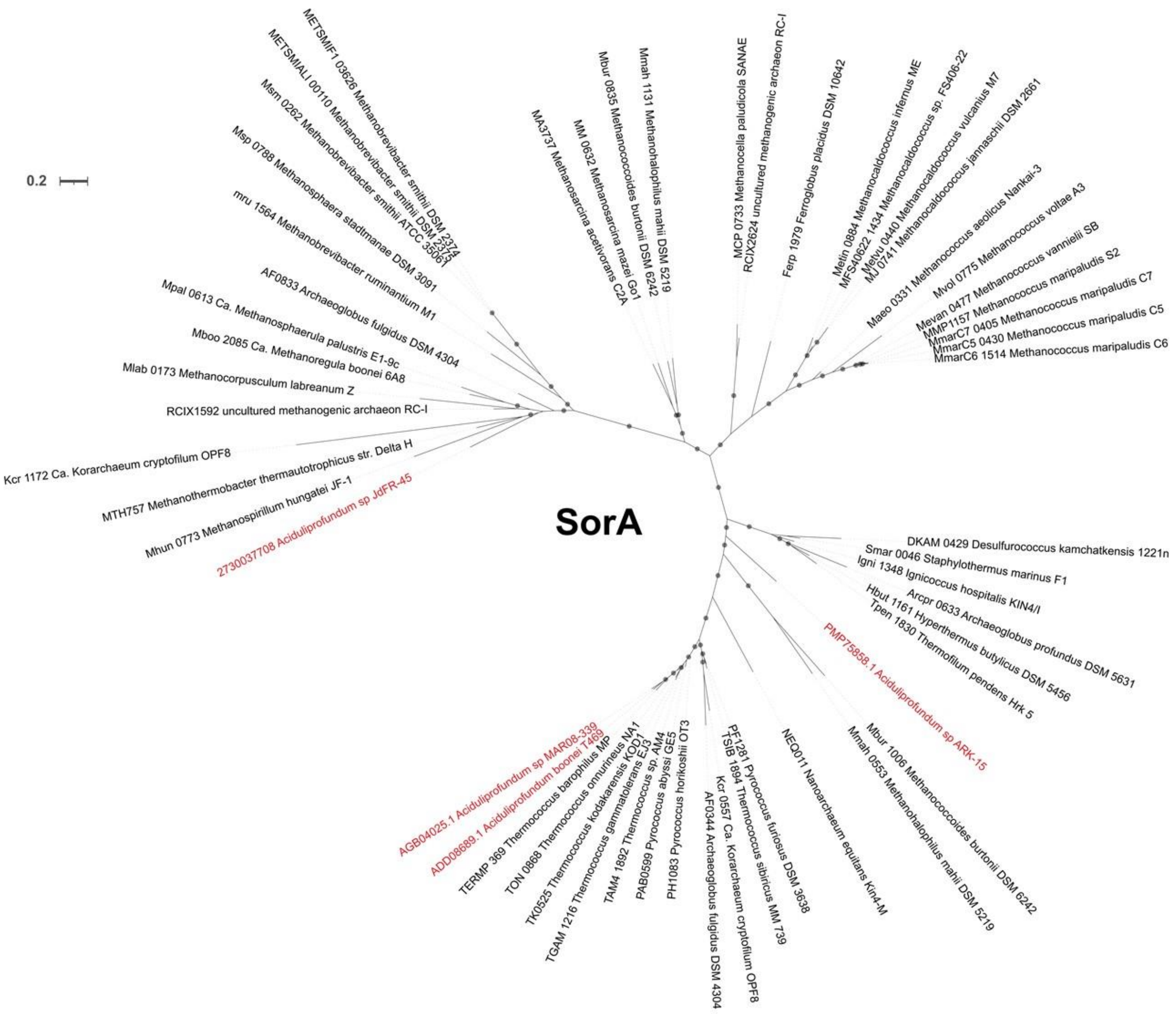


**Figure S2.** Unrooted phylogenetic tree constructed from the predicted amino acid sequences of SorA in *Aciduliprofundum* and Archaeal sequences from sorGOdb. In red color are sequences from *Aciduliprofundum.* Bootstrap values over 60 are represented with gray circles in the respective branches. Scale bar represent 0.2 amino acid substitutions per site.

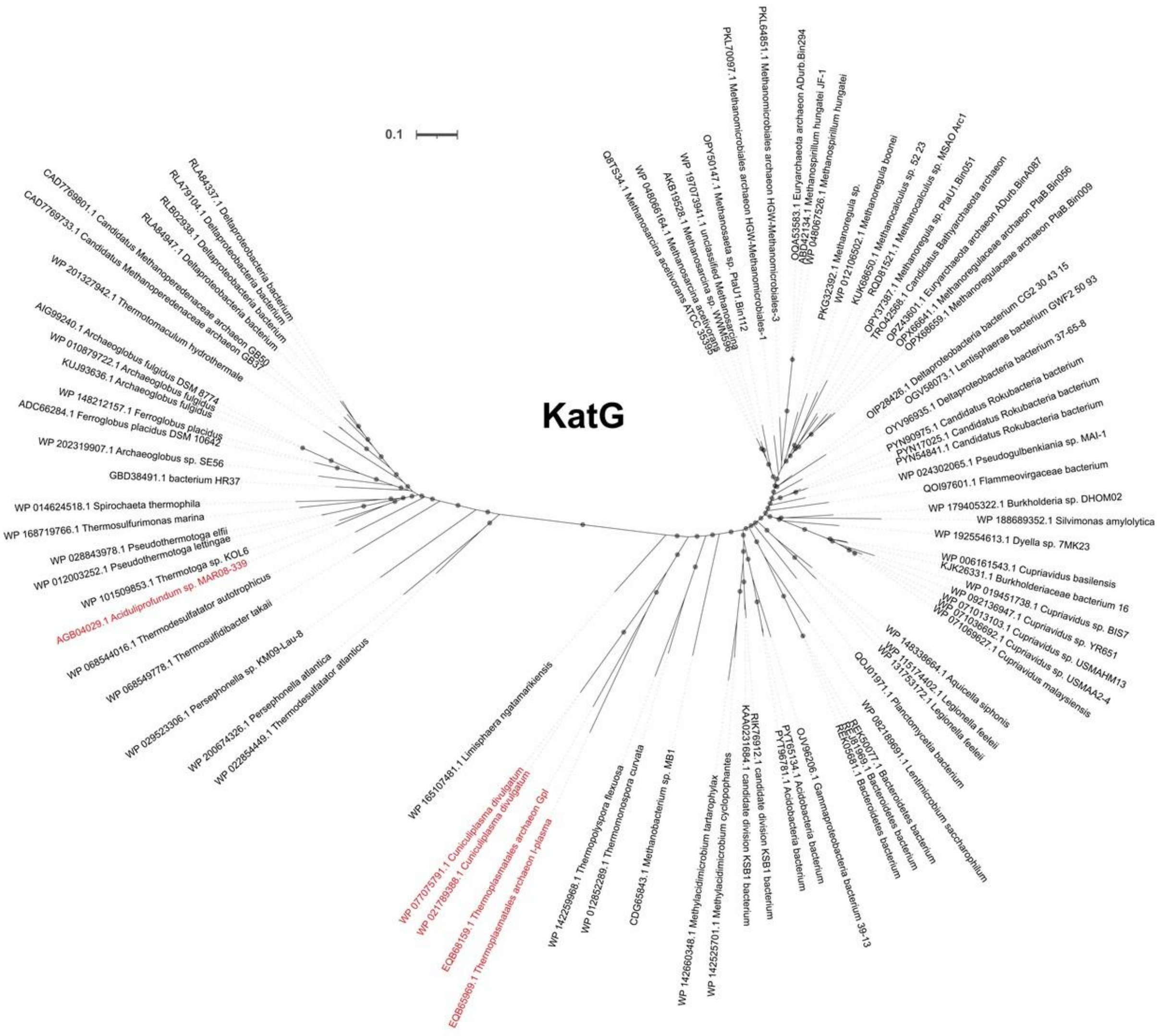


Figure S3: Unrooted phylogenetic tree constructed from the predicted amino acid sequences of KatG in *Aciduliprofundum, Cuniculiplasma* and *Thermoplasmatales* and their best hits in the NCBI. In red color are sequences from *Aciduliprofundum.* Bootstrap values over 60 are represented with gray circles in the respective branches. Scale bar represent 0.1 amino acid substitutions per site.

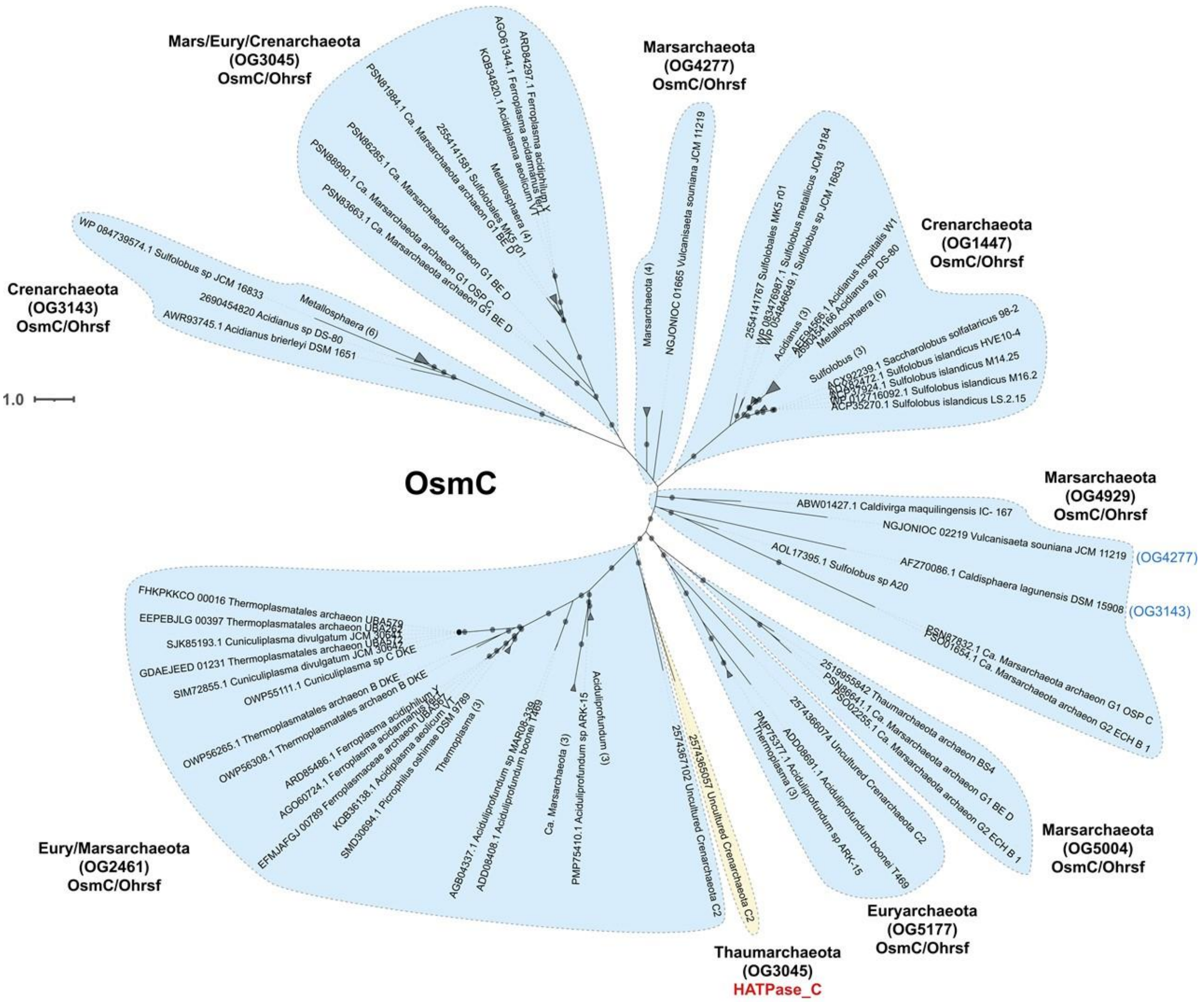


**Figure S4** Unrooted phylogenetic tree constructed from the predicted amino acid sequences of OsmC in the different acidophilic Archaea. Each clade is delimited by a blue background. The specific phyla with sequences in each clade are annotated with the corresponding orthogroup. In yellow is shown the sequence of Thaumarchaeota that doesn't share the main functional annotation of OsmC of the other sequences Bootstrap values over 60 are represented with gray circles in the respective branches. Scale bar represent 0.1 amino acid substitutions per site.

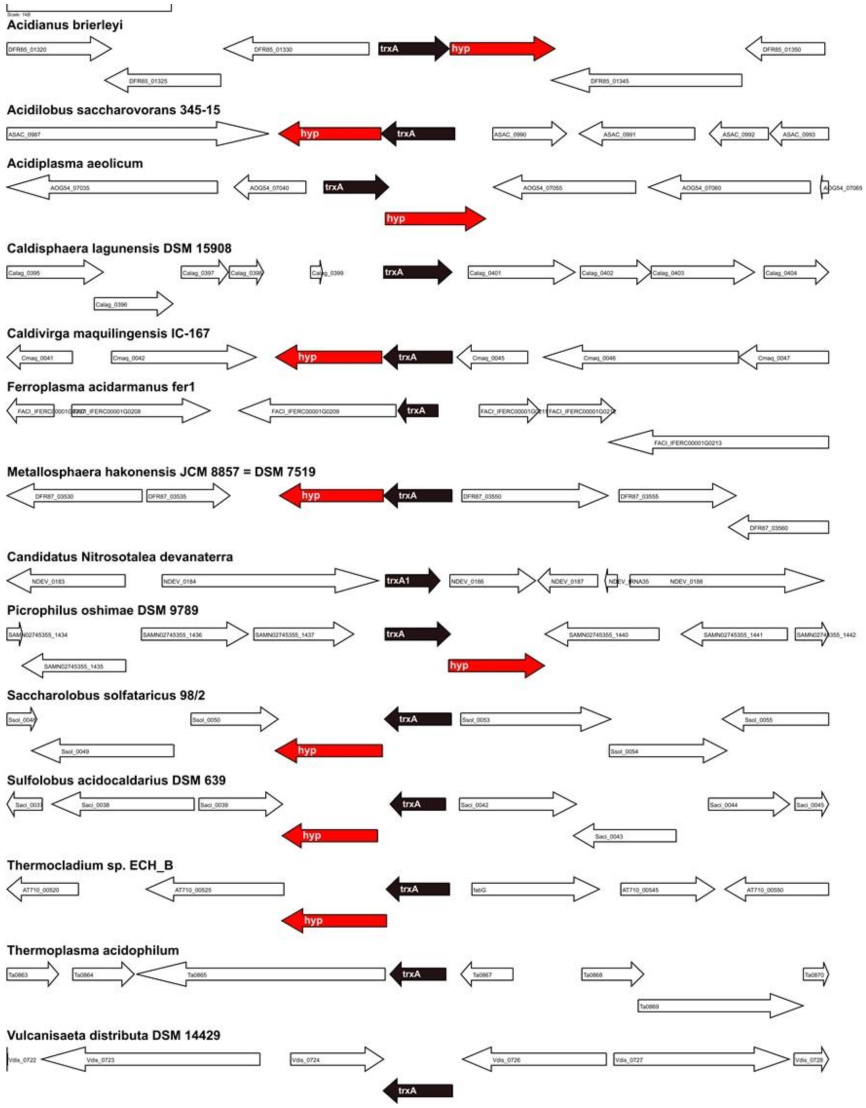


**Figure S5.** Genomic context of *trxA* and the conserved hypothetical protein OG0923 in several representative genomes. Locus tag for genes are shown inside each of the arrows. Color Coding: red=hypothetical OG0923, black = *trxA*. White: non-conserved genes
